## Supplemental _Material for "Decoding functional high-density lipoprotein particle surfaceome interactions"

### Supplementary Tables:

**Table S1**: List of identified peptides in auto-CSC experiments with EA.hy926 cells, HAECs, HEPG2 cells, THP1 cells, activated THP1 cells, and foam cells. Identifications are filtered for FDR < 0.01 and NxS/T motifs.

**Table S2**: List of ranked and scaled protein abundances and the peptide counts captured in auto-CSC experiments with EA.hy926 cells, HAECs, HEPG2 cells, THP1 cells, activated THP1 cells, and foam cells.

**Table S3**: Matrix of all identified proteins in auto-CSC experiments with EA.hy926 cells, HAECs, HEPG2 cells, THP1 cells, activated THP1 cells, and foam cells.

**Table S4**: Safequant output of the auto-CSC experiments quantitatively comparing VEGF-A treated vs untreated HAEC surfaceomes.

**Table S5**: Statistics from the gene ontology analysis of genes regulated by VEGF-A in HAECs.

**Table S6**: Gene ontology protein annotation of the analysis of VEGF-A-sensitive proteins on HAECs.

**Table S7**: Protein content of rHDL as identified by Comet. Peptide and protein identifications were filtered with an FDR of 1%. Common contaminants and keratins were removed from the protein list.

**Table S8**: Safequant output of the HATRIC-LRC experiment with APOA1, lipidated APOA1, and HDL as ligands on EA.hy926 cells.

### Supplementary figures:

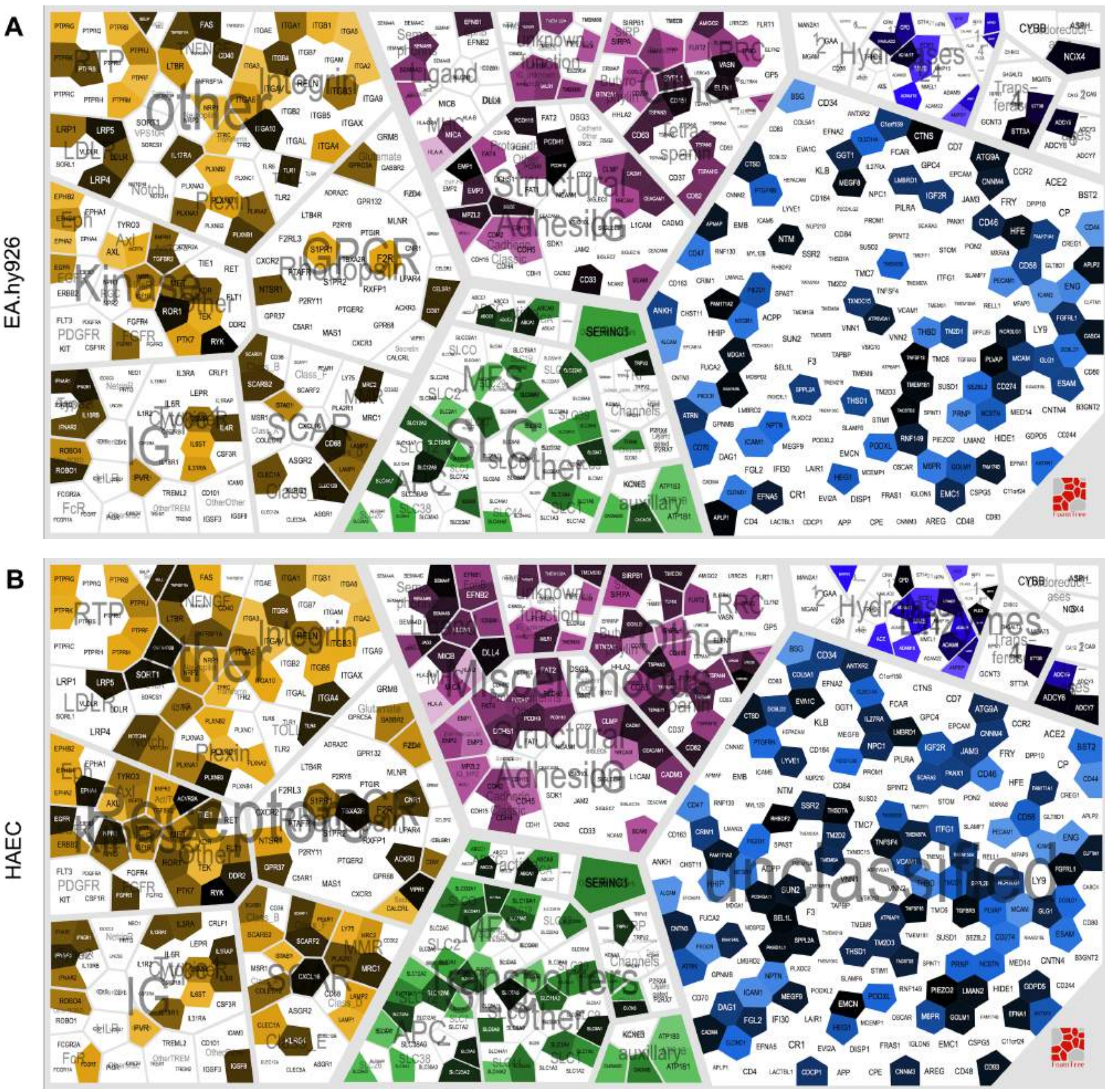

**C**

HEPG2

**D**

THP1

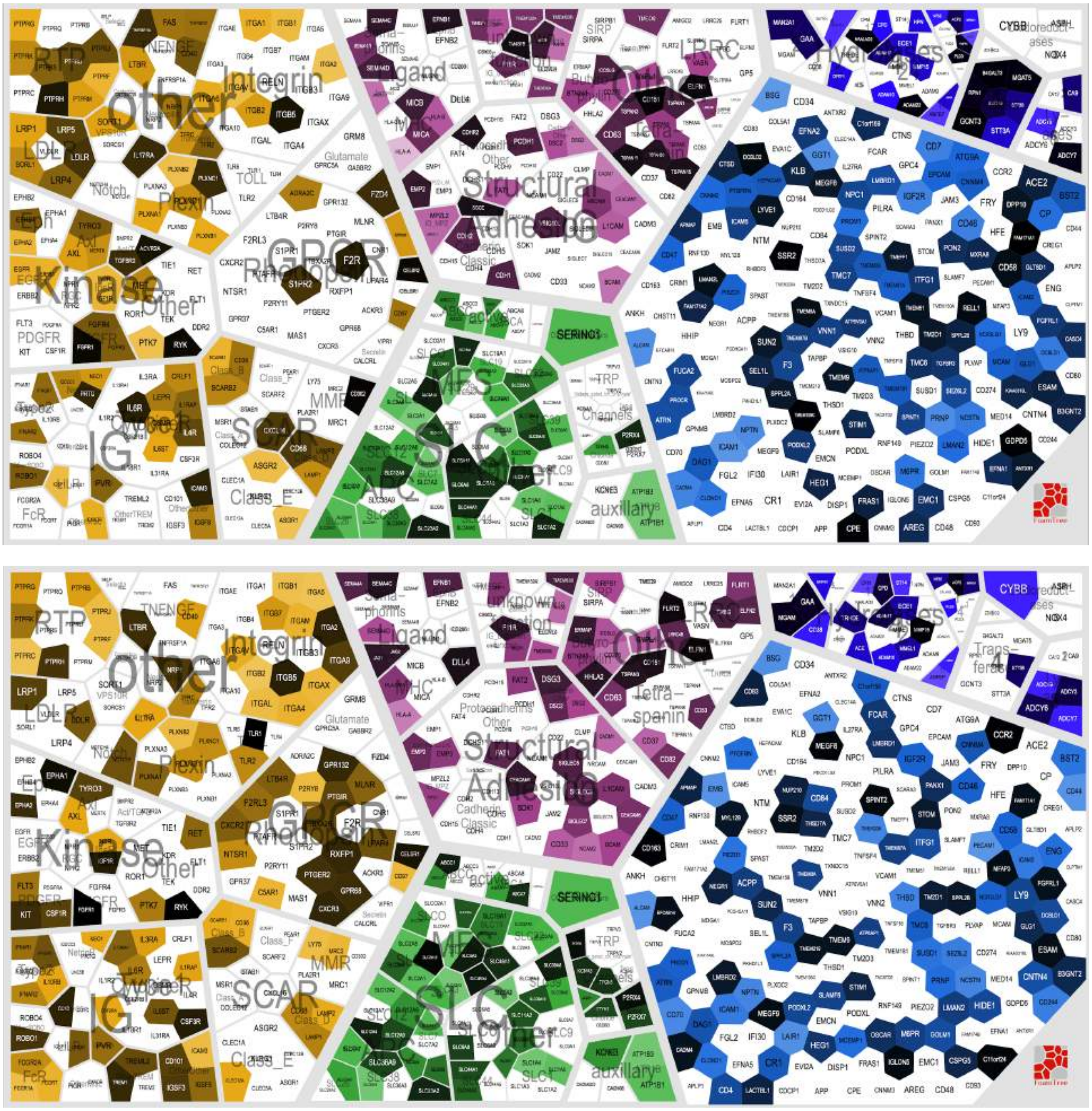

**E**

activated THP1

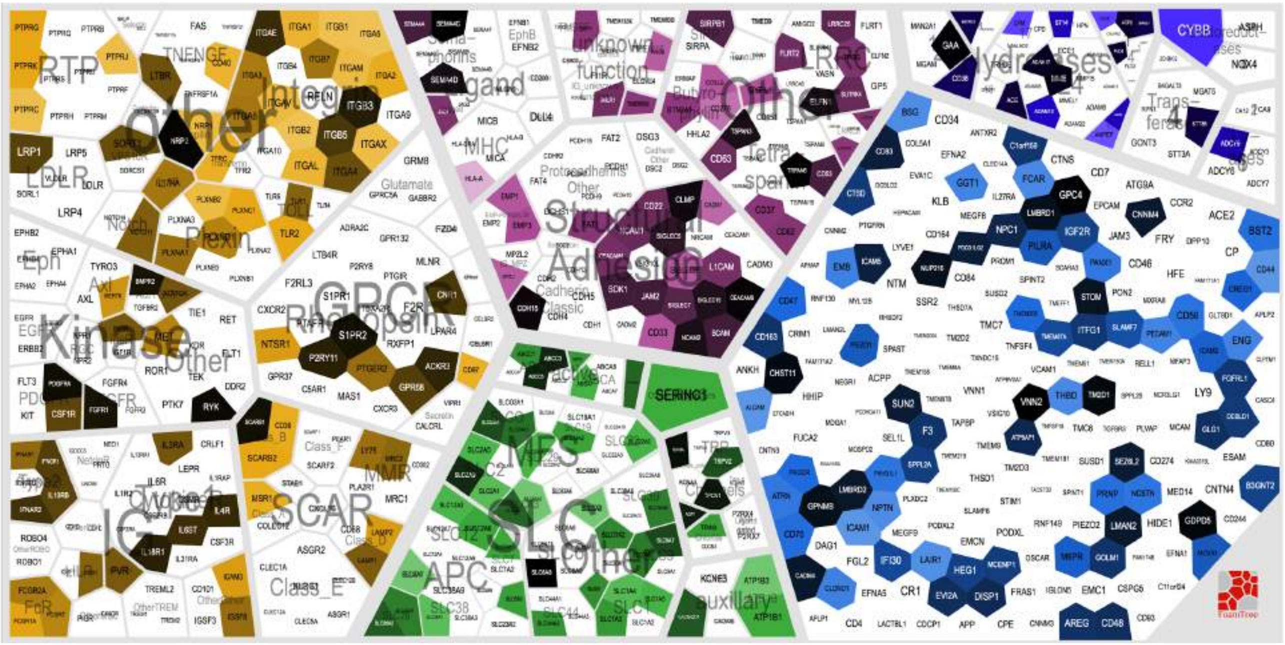

**F**

Foam cells

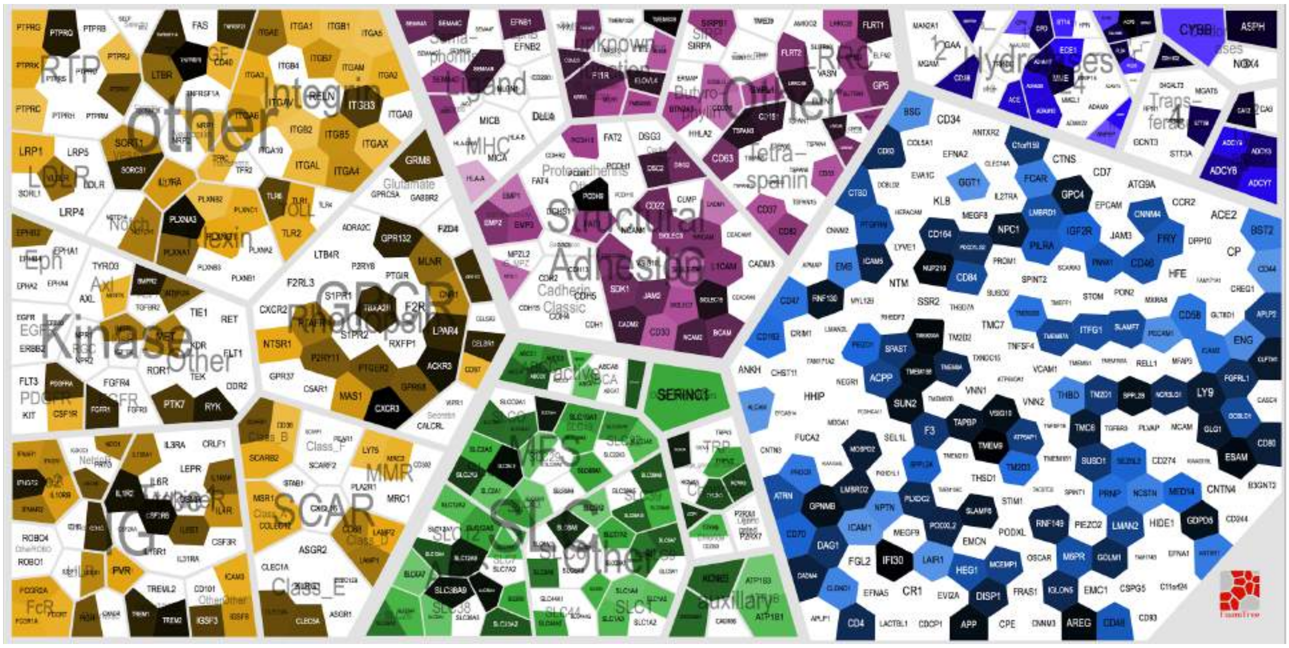

**Fig. S1:** Voronoi treemaps of the quantitative surfaceome of different cell lines. **A)** EA.hy926 cells, **B)** HAECs, **C)** HEPG2 cells, **D)** THP1 cells, **E)** activated THP1 cells, and **F)** foam cells. The background proteome comprises all identified proteins across the different cell types. Color scale corresponds to the cell type-specific ranked and scaled abundances of the proteins.

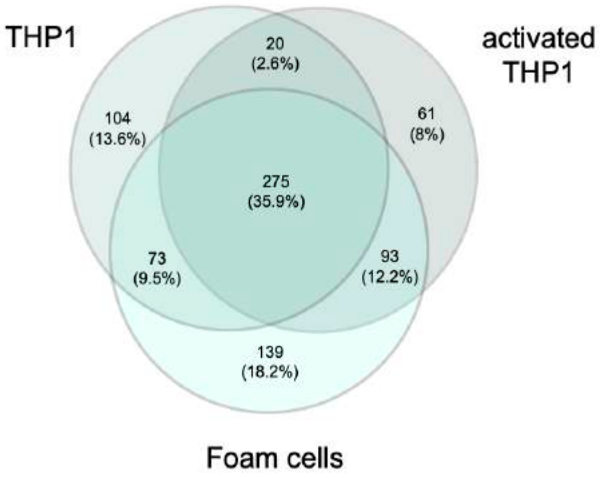


**Fig. S2**: Venn diagram of overlap of identified proteins in the auto-CSC experiments with THP1 cells, activated THP1 cells, and foam cells.

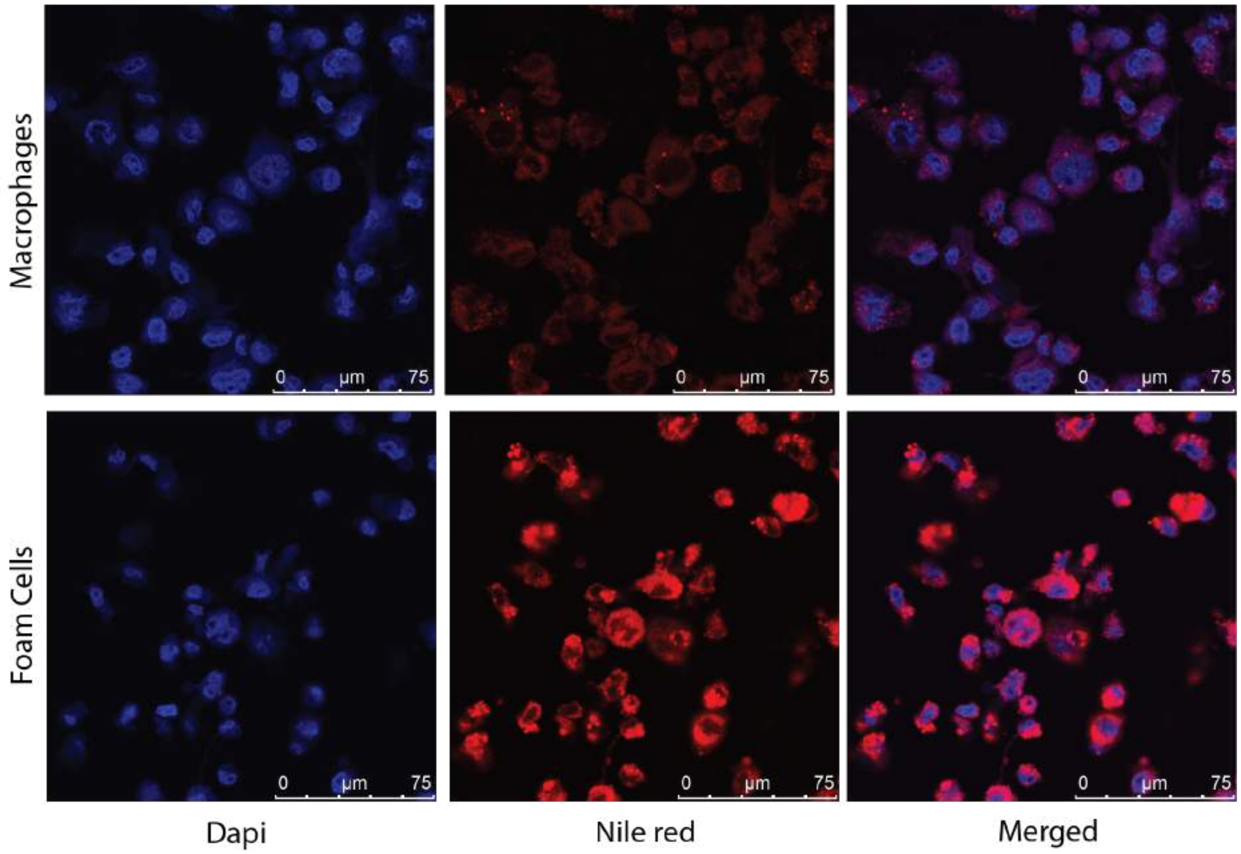


**Fig. S3:** Microscopy images of the lipid content within THP1 cells, activated THP1 cells, and foam cells. Lipids were stained with Nile red.

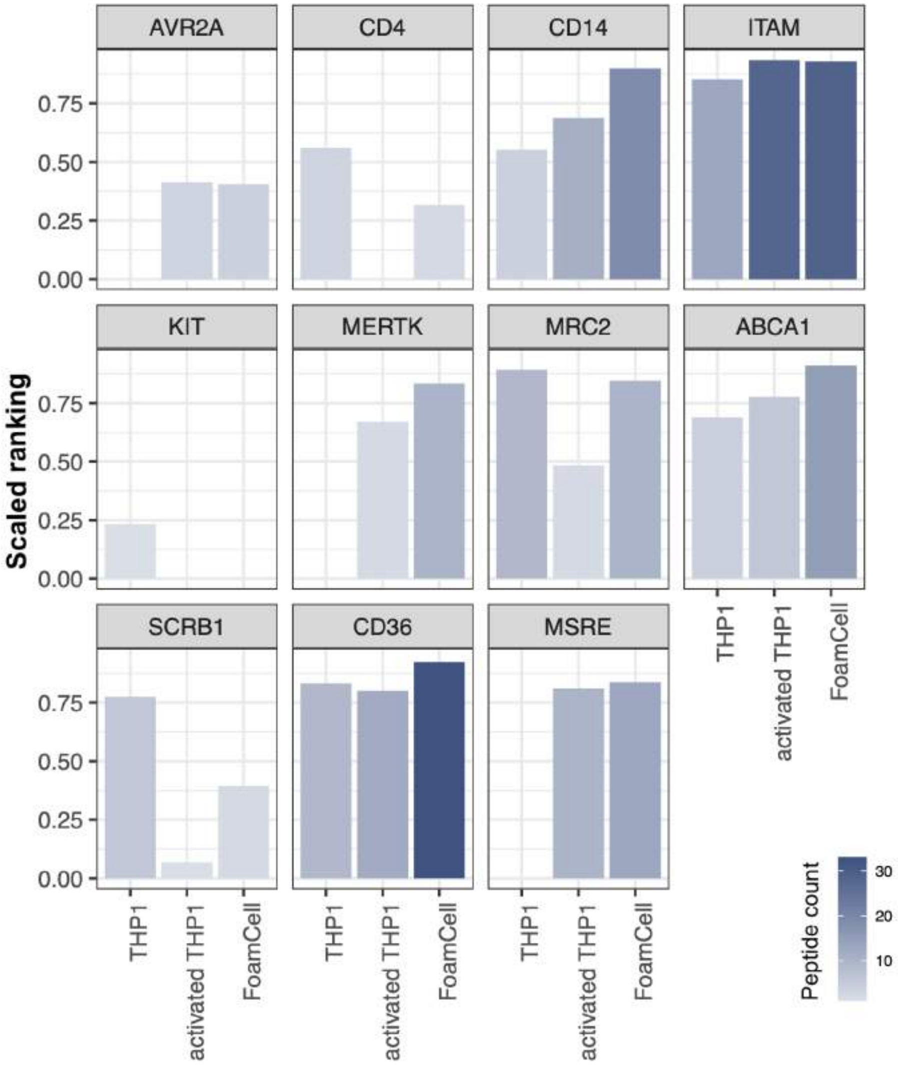


**Fig. S4**: Bar charts of selected proteins and their scaled ranked abundances determined with auto-CSC on THP1 cells, activated THP1 cells, and foam cells. Included are proteins previously shown to have altered surface abundance upon THP1 differentiation (1): AVR2A, CD4, CD14, ITAM, KIT, MERTK, and MRC2. Data for the APOA1/HDL receptors ABCA1 and SCRB1 as well as the oxLDL receptors CD36 and MSRE (2) are also shown.

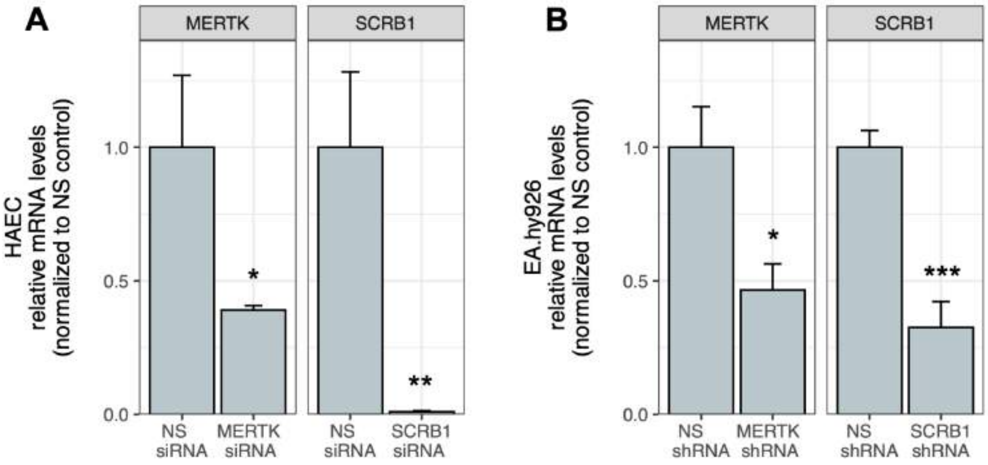


**Fig. S5:** mRNA levels upon silencing of *MERTK* and *SCRB1* in EA.hy926 and HAECs. Significance of *MERTK* and *SCRB1* silencing using shRNA in EA.hy926 cells and siRNA in HAECs compared to control cells treated with non-silencing (NS) vector or control, respectively, was assessed with the Student's t-test (* $p < 0.05$, ** $p < 0.01$, *** $p < 0.001$).

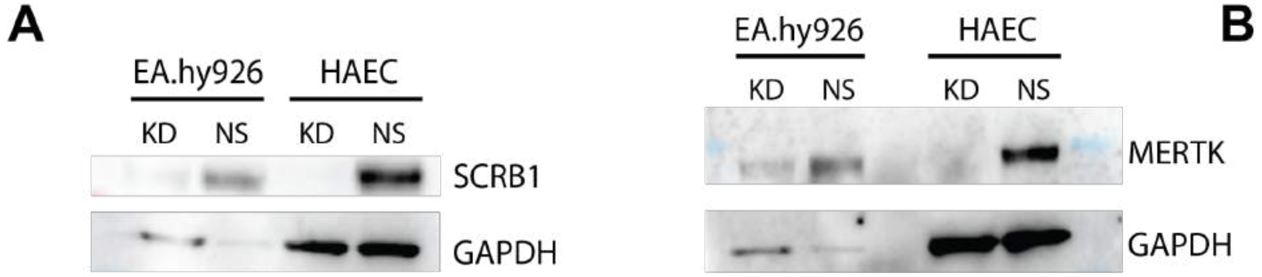


**Fig. S6:** Western blot analysis of total protein abundances upon silencing of **A)** *SCRB1* and **B)** *MERTK* in EA.hy926 cells and HAEC with shRNA or siRNA, respectively (KD), and upon treatment with non-silencing controls (NS).

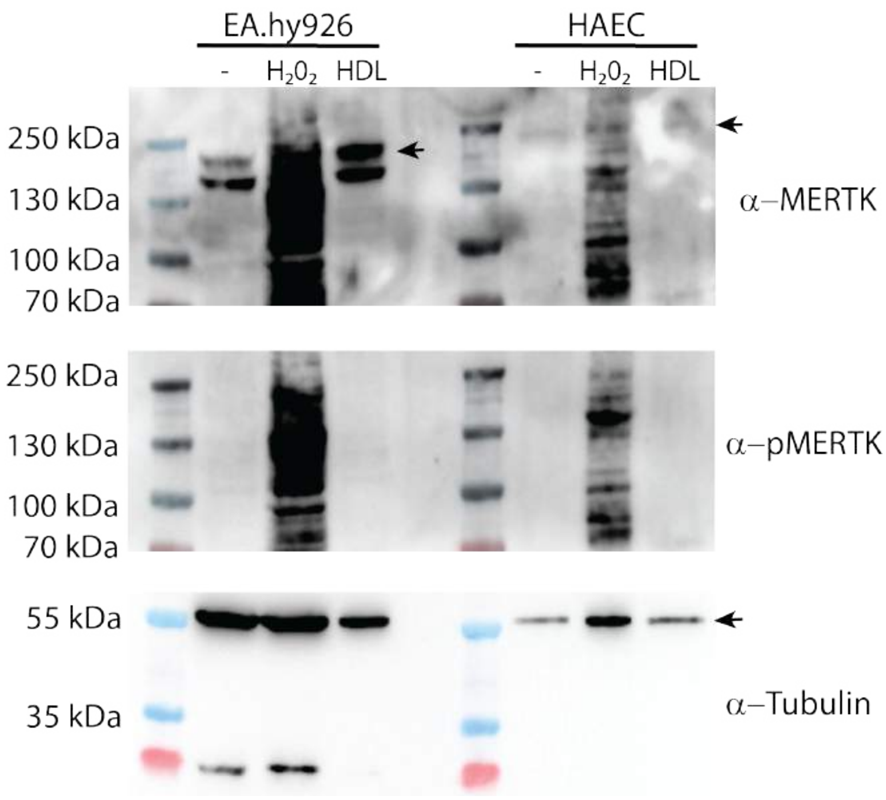


**Fig. S7:** Effect of HDL treatment on phosphorylation of MERTK in EA.hy926 cells and HAECs as shown by western blot. $H_2O_2$ treatment served as positive control (3). Tubulin was used as loading control.
